# Construction of mirror-image RNA nanostructure for enhanced biostability and drug delivery efficiency

**DOI:** 10.1101/2024.07.17.603907

**Authors:** Ying Zhang, Yuliya Dantsu, Wen Zhang

**Affiliations:** Department of Biochemistry and Molecular Biology, Indiana University School of Medicine, 635 Barnhill Drive, Indianapolis, IN 46202; Melvin and Bren Simon Comprehensive Cancer Center, 535 Barnhill Drive, Indianapolis, IN 46202, USA

**Keywords:** mirror-image RNA, chemotherapy, siRNA, folic acid

## Abstract

The development of stable and efficient drug delivery systems is crucial for advancing therapeutic applications. Here, we present a novel approach involving the construction of a mirror-image RNA (L-RNA) nanostructure to enhance biostability and drug delivery efficiency. Specifically, we utilized L-RNA to create a three-way junction structure, which was then conjugated with small interfering RNA (siRNA) and chemotherapy agents for targeted drug delivery. Our findings demonstrate that this L-RNA nanostructure significantly improves therapeutic efficiency due to its enhanced stability compared to natural D-RNA. In additionally, the conjugated folic acid (FA) group dramatically enhance the specificity and endosomal escape efficiency of the nanoparticles, which benefit the combinatorial drug delivery. This work highlights the potential of mirror-image RNA nanostructures as robust platforms for drug delivery applications.

## INTRODUCTION

The development of efficient and targeted drug delivery systems is critical in modern medicine, particularly for the delivery of small interfering RNA (siRNA) and chemotherapeutic agents. Traditional drug delivery methods often suffer from significant drawbacks, including poor targeting, rapid clearance from the bloodstream, and degradation by enzymatic activity, which collectively reduce therapeutic efficacy and increase the risk of off-target effects and toxicity. Lipid nanoparticles, viral vectors, and polymer-based systems have been employed to address some of these challenges, but they frequently face issues such as immunogenicity, limited loading capacity, and complex manufacturing processes.

In recent years, RNA nanotechnology has emerged as a promising alternative for drug delivery, leveraging the programmable nature of RNA to create versatile and multifunctional nanostructures. RNA, as a simple linear polymer, is composed of merely four types of nucleobases, exterior phosphate groups, and a puckering backbone of furanose^1^. In addition to its therapeutic potential, RNA possesses inherent properties, making it a promising material for constructing nanostructures for drug delivery^2, 3^. Such properties include high thermodynamic stability, the capacity to fold into a large number of distinct structures, and its nanoscale size, which enhances its permeability in vivo^4^. Hence, these unique characteristics make RNA an ideal candidate to use in novel drug delivery systems. In laboratories, RNA nanostructures have been precisely designed to achieve desired shapes and functions, enabling the creation of complex architectures that can carry multiple therapeutic agents. These RNA-based systems benefit from biocompatibility and the ability to interact specifically with cellular targets through Watson-Crick base pairing.

RNA nanotechnology offers tremendous potential in disease treatment, nevertheless, it is hindered by their natural susceptibility to degradation by nucleases, which drastically reduces their stability and limits their practical use^5^. The abundance of RNase enzymes in biological systems renders RNA nanoparticles highly vulnerable to degradation, resulting in a short half-life of merely 5 to 10 hours in vivo^6, 7^. Presently, the predominant strategy towards solving the stability issue of RNA therapy involves the incorporation of sugar modifications, such as 2′-F or 2′-OMe^8^. However, while these chemical derivatizations can extend the circulation time of RNA in vivo to a certain degree, their efficacy is limited^9, 10^. Moreover, these sugar modifications may potentially give rise to unanticipated toxicities^11, 12^ and perturbation of the native RNA conformation^13^. As such, the exploration of innovative RNA materials is crucial for the development of RNA-based therapeutics.

Mirror-image nucleic acids, also known as L-RNA or Spiegelmers, offer a robust solution to the stability issue faced by traditional RNA molecules. L-RNA consists of nucleotides that are mirror images of the naturally occurring D-nucleotides^14^. This chirality prevents recognition and degradation by natural ribonucleases, conferring exceptional stability in biological environments^15, 16^. Similar to D-RNA, L-RNA oligonucleotides can be chemically modified, thereby broadening their scope of functionalities^17-20^. These features make L-RNA uniquely suited to help to address some long-standing bottlenecks in the field of nucleic acid drug development. Successful examples of using L-nucleic acid in therapeutic applications include the use of L-nucleoside molecules as antiviral and antitumor agents^16^, L-RNA aptamers to bind to disease targets^21-24^, and the of L-DNA tetrahedron structures as drug delivery vehicles^25^. Given the advantages, the use of L-RNA in constructing nanostructures thus presents a significant advancement in the field of RNA nanotechnology, combining the structural and functional benefits of RNA with enhanced biostability.

In this study, we have developed a novel L-RNA-based nanostructure to enhance drug delivery efficiency and stability. By constructing a three-way-junction (3WJ) structure using L-RNA, we create a stable platform capable of conjugating both siRNA and chemotherapeutic agents. This innovative approach aims to exploit the inherent stability of L-RNA to improve the therapeutic efficiency of the conjugated drugs. Our primary objectives are to evaluate the biostability of the L-RNA nanostructure in comparison to traditional D-RNA-based systems and to assess its therapeutic efficacy in delivering siRNA and chemotherapy agents. We hypothesize that the enhanced stability of the L-RNA nanostructure will lead to increased bioavailability and therapeutic efficacy of the delivered agents, providing a significant improvement over existing RNA-based delivery systems.

This work represents a critical advancement in the application of mirror-image RNA in drug delivery. By demonstrating the potential of L-RNA nanostructures to enhance drug stability and therapeutic outcomes, our study paves the way for the development of next-generation delivery platforms that could revolutionize the treatment of various diseases, including cancer and genetic disorders.

## MATERIALS AND METHODS

### Preparation of L-RNA and L-RNA/L-DNA chimeric oligonucleotides, and functional L-RNA three-way-junction (3WJ) nanoparticles

The L-RNA, and chimeric L-RNA/L-DNA oligonucleotides used for biochemical and cellular studies were solid-phase synthesized using ABI 394 DNA/RNA synthesizer. The phosphoramidite compounds are from Chemgenes or Glen Research, Inc. The DMTr-on in-house synthesized oligonucleotides were deprotected by AMA (1:1 mixture of ammonium hydroxide and methyl amine solution) for 15 min at 65°C, followed by the Et_3_N•3HF treatment; and purified by Agilent ZORBAX Eclipse-XDB C18 column using 25 mM triethylammonium acetate in H_2_O (pH 7.5) with an increasing gradient of 0 % to 30 % acetonitrile over 40 mins. The 5’-Cyanine 5 labelled oligonucleotides were deprotected by conc. ammonium hydroxide at room temperature for 36 hours, followed by the Et_3_N•3HF treatment and denaturing urea polyacrylamide gel electrophoresis. To label the L-RNA with folic acid (FA), the alkyne modified L-RNA was synthesized using the 1-Ethynyl-dSpacer CE Phosphoramidite from Hongene Biotech, Inc., following the standard deprotection protocol. After purification, the oligonucleotides were desalted and re-concentrated to the proper concentrations for nanoparticle construction and biochemical experiments.

The Cy5 labelled, FA labelled, and wild-type L-3WJ nanoparticles are composed of three oligonucleotides. The L-3WJ nanoparticle containing siRNA is composed of four oligonucleotides. All the sequence information is listed in Supporting Information. All the nanoparticles were self-assembled by mixing the different strands at the same molar concentrations in TMS buffer (25 mM Tris-HCl pH 7.5, 10 mM MgCl_2_, 50 mM NaCl) or PBS buffer (137 mM NaCl, 2.7 mM KCl, 10 mM Na_2_HPO_4_, 2 mM KH_2_PO_4_, pH 7.4), heated to 90°C for 2 min and slowly cooled to 23 °C. After the annealing program, the nanoparticles were loaded onto 10% native PAGE, stained by Sybr Gold (Invitrogen), and visualized by Bio-rad ChemiDoc Imaging System.

### Click reaction to label L-RNA with folic acid molecule

To facilitate the specific cellular targeting, the L-RNA was synthesized with alkyne modification, which later was conjugated with a folic acid molecule via a click reaction. The folic acid-PEG-azido compound was purchased from Biopharm PEG, Inc. To the alkene modified L-RNA solution, excess of folic acid-PEG-azide, tris(3-hydroxypropyltriazolylmethyl)amine/CuSO_4_ and sodium ascorbate solution were added. The reaction continued for 30 min, followed by ethanol precipitation and gel purification, to get folic acid labelled L-RNA. Details of the click reaction procedure are included in the Supporting Information.

### L-RNA enzymatic stability assays

Both L- and D-3WJ nanoparticles were incubated in 25% Human serum off-the-clot (Sigma) at 37°C for different times. 200 ng of RNA were taken at 30 min, 1 hr, 2 hr, 8 hr and 24 hr time points, and analyzed by 10% native PAGE in TB buffer. The gels were stained with Sybr Gold (Invitrogen) and washed briefly before imaging using Bio-rad ChemiDoc Imaging System.

### Melting temperature measurement of L-RNA nanoparticle

Melting temperature measurement was performed using the QuantStudio Real-time qPCR equipped with temperature controller system. 1× SYBR Green I dye (Invitrogen, emission 465-510 nm) was added to the appropriate buffer (750 mM NaCl, 10 mM MgCl_2_, 50mM Na_3_PO_4_, pH 6.8) of different RNA nanoparticles (1 µM concentration), and the fluorescence signals were monitored. The nucleic acid samples were heated to 90°C for 2 min, then slowly cooled down to 4°C at the ramping rate of 0.1°C/s. The raw data were analyzed by QuantStudio software using the first derivative of the melting profile. The *T*_m_ value represents the mean and standard deviation of three independent experiments.

### Colony-forming unit (CFU) assay

The toxicity of L-RNA nanoparticles to bone marrow progenitor cells was performed in the In Vivo Therapeutics Core at the Indiana University Simon Comprehensive Cancer Center^26^. Colony Forming Unit (CFU) assay was performed using enriched human CD34^+^ Hematopoietic stem and progenitor cells (HSPCs) cultured in Methocult GF (Stem Cell Technologies, Inc., Cat. # H4434). The cells were seeded in triplicate dishes at concentrations of 2 × 10^3^ to obtain 60-70 colonies per 35 mm dish. For survival assays, cells were exposed to L-RNA 3WJ nanoparticles at different concentrations prior to plating. After 14 days of incubation at 37°C in 5% CO_2_, colony forming units were enumerated using the Axiovert 25 inverted light microscope.

### Cell culture

MDA-MB-468 (RRID: CVCL_0419) and HEK-293T (RRID: CVCL_0063) cells were purchased from the American Tissue Type Collection (ATCC, Manassas, VA), or gift from Indiana University Melvin and Bren Simon Comprehensive Cancer Center. Cells were maintained in DMEM (Corning, Cat. #10-013-CV) supplemented with 10% FBS. All cell lines were maintained at 37°C in a fully humidified atmosphere containing 5% CO_2_, used at early passage for experiments, and tested to be free of mycoplasma contamination.

### Western blot for MCL1 gene expression in different cells

Cells were lysed using 1% SDS extraction buffer supplemented with protease inhibitors (Santa Cruz Biotechnology, TX, USA). Briefly, the cell extract was heated at 95°C for 5 minutes and sonicated (4 pulses, 4 cycles). Denatured samples (20–40 μg) were subjected to SDS-PAGE, and proteins were transferred onto nitrocellulose membranes by electrophoretic transfer. Non-specific binding sites were blocked at room temperature for 1 hour with 5% (w/v) Blotting-Grade milk (Bio-Rad Laboratories, CA, USA) in Tris-buffer saline (Boston Bio Products, MA, USA) containing 0.05% (v/v) Tween-20 (Thermo Fisher, MA, USA) (TBS-T). The membranes were incubated overnight with the Anti-MCL1 antibodies (Santa Cruz Biotechnology, TX, USA) and then with the peroxidase-conjugated secondary antibody for 1 hour. The signal was captured using a Bio-Rad ChemiDoc imager, and the band intensities were analyzed by densitometry on Image Lab (Bio-Rad Laboratories, CA, USA) or ImageJ software. Analysis of beta-actin expression is used as control.

### Confocal microscopy imaging of internalization of L-RNA nanoparticles

Cells were grown on poly-d-lysine-coated glass cover slides in normal growth media as described above and allowed to attach overnight. Cells were washed once with Opti-MEM medium (Thermo, Cat. #11058021) prior to treatment with Cy5- and FA-labelled L-3WJ nanoparticles diluted in Opti-MEM medium (Thermo, Cat. #11058021).

For confocal microscopy, cells were fixed by 4% paraformaldehyde, permeabilized using PBS-T (0.01% Triton X) and stained by DAPI for nuclei visualization. FITC-conjugated phalloidin is used to stain the cell cytoskeleton. Confocal microscopy was performed using a Zeiss AxioObserverZ1 modified by 3i for confocal microscopy with the addition of a CSU-X1 M1 Spinning Disk Confocal and a Prime BSI CMOS Detector.

### Formation of DOX/nanoparticle complexes

Doxorubicin hydrochloride (HPLC-purified, Sigma-Aldrich) was dissolved in Milli-Q water for a 2 mM stock solution, divided into aliquots and stored at −20°C. DNase I to treat L-nanoparticle was purchased from Sigma-Aldrich. A stock solution was prepared at 2 U/ul concentration in deionized water and stored in aliquots at −20°C, and after thawing stored at room temperature (RT) and used within the same day.

Doxorubicin molecule was loaded onto the double-stranded L-DNA by annealing the DOX/L-DNA complex (at appropriate ratios) and storing the solution at room temperature for 24 h. The DOX^+^ L-3WJ nanoparticle was formed by mixing the DOX/L-DNA complex with FA-L-3WJ containing L-DNA overhang at 1:1 ratio for 24 h. The assembled FA-DOX^+^ L-3WJ nanoparticle was used for cellular experiments with the corresponding concentrations.

Spectra of L-DNA and its DOX complexes were recorded at pH 7.4 with SpectraMax Plus 384 spectropolarimeter in a 10 mm optical path length cuvette, in the Chemical Genomics Core, IUSM. The fluorescence spectra were recorded at λ_ex_ = 480 nm and λ_em_ from 500 to 700 nm. All measurements were performed at room temperature.

### Alamar blue assay for cell toxicity measurement

For Alamar blue assay, cells were seeded into poly-d-lysine-coated 96-well plates at 8000-10000 cells per well in normal growth media as described above and allowed to attach overnight. Different drugs were diluted in Opti-MEM medium (Thermo, Cat. #11058021) to appropriate concentrations before the treatment. Cells were washed once with Opti-MEM medium (Thermo, Cat. #11058021) prior to treatment with different drugs. After treatment, Alamar Blue reagent (Invitrogen, Cat. #A50101) was added to each well, with the final volume of 10% in the well. The cells with Alamar Blue were further incubated at 37°C for 6 hours, and the fluorescence intensity (excitation 560 nm, emission 590 nm) was measured using Synergy H1 microplate reader (BioTek Instruments, Inc., VA). The fluorescence readings were analysed to determine cell viability, and the data were plotted.

## RESULTS

### Design, construction and characterization of L-RNA nanoparticle

An array of RNA junctional structures have been identified across various species in nature, possessing the unique ability to fold into diverse nanostructures, each with the distinct shapes and stabilities^27, 28^. Here we choose the packaging RNA (pRNA) of bacteriophage phi29 DNA to construct our L-RNA three-way-junction (L-3WJ) nanoparticle (Figure 1B)^29^. The native 3WJ structure from bacteriophage phi29 was assembled using three separate pieces of RNA, and the resulting nanoparticle shows resistance to denaturing reagent and high temperature^29^. We utilized these same sequences of native pRNA, and constructed the L-3WJ nanoparticle composed of L-RNAs (L-3WJ_a_, L-3WJ_b_, L-3WJ_c_), each present in equal molar concentrations. The formation of the L-3WJ was confirmed by 10% native PAGE (Figure 1C). A stepwise assembly between the L-RNA oligonucleotides was observed, including the formation of dimers and the assembled trimer, in high yield. Our data agrees with the reported assembly of native RNA three-way-junction nanoparticle^30^. In addition, L-3WJ RNA strands don’t form 3WJ structures with D-RNAs, indicating the inability to form Watson–Crick base paired structures between D- and L-oligonucleotides.

**Figure 1.**
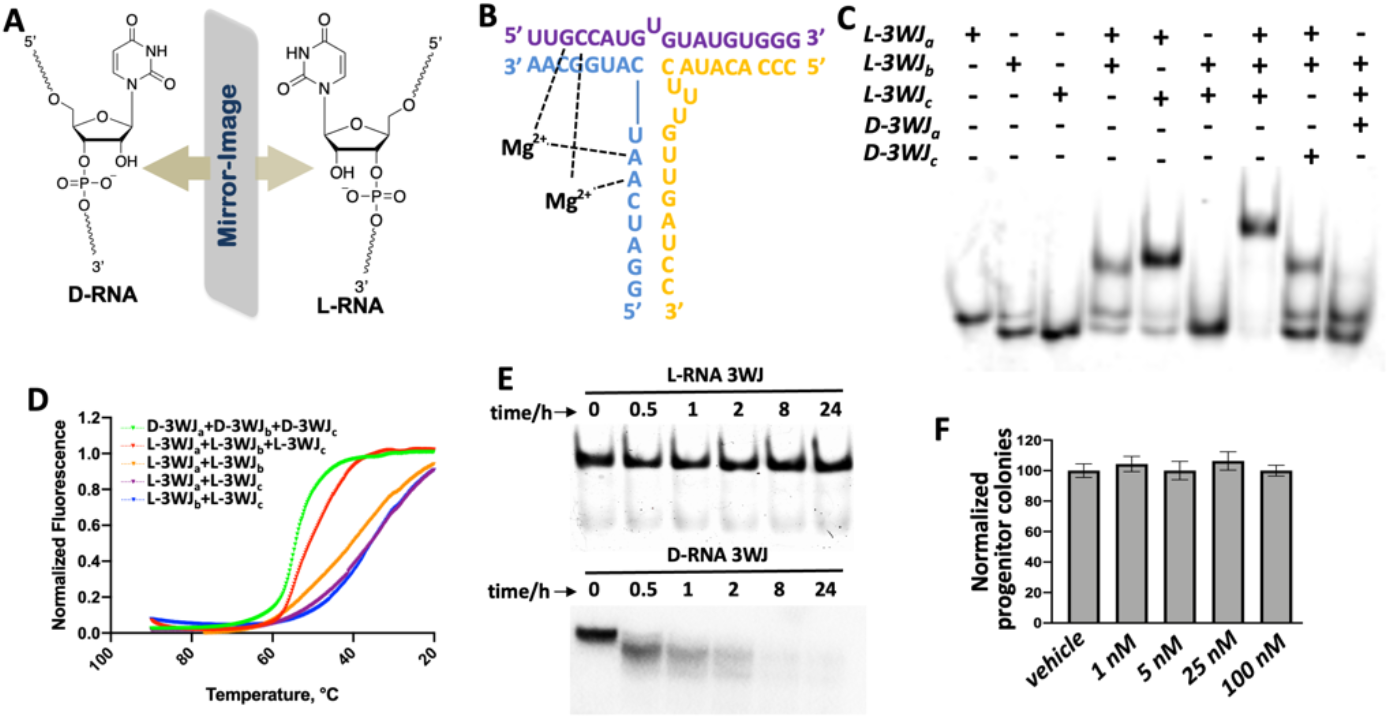
Design, Construction and Characterization of L-RNA 3WJ Nanoparticle. (A) Chemical structures of D- and L-ribonucleotides; (B) 3WJ domain composed of three L-RNA oligomers in blue, yellow and purple. (C) 10% native PAGE showing the assembly of the L-3WJ core, stained by SYBR gold. (D) *T*_m_ melting curves for the assembly of the L-3WJ core. Melting curves for the L-RNA two-strand combinations (blue, yellow, purple), the L-RNA three-strand combination (red) and D-RNA three-strand combination (green) are shown. (E) Stability assay of L-3WJ (top) and D-3WJ (bottom) after treating with serum. (F) The colony-forming units from vehicle and L-RNA 3WJ treated groups were analyzed. Error bars represent the mean ± SEM from three independent experiments (n = 3).

### Stability of L-3WJ nanoparticle

The thermal stability of the L-3WJ nanoparticles were determined by measuring the fluorescence intensities as a function of temperature in the presence of SYBR Green II dye. The result indicates the annealing transition of different complexes of RNA assemblies (Figure 1D). The nanoparticles of three strands, including D-RNA and L-RNA, produced the highest annealing temperature compared to the dimers. The melting temperature of L-3WJ was measured to be 57.3 °C, which is comparable to the melting temperature of D-3WJ (T_m_ is 59.8 °C). This similar thermostabilities are due to the same physical and chemical properties of mirror-image oligonucleotides, and the slight difference is likely from the different qualities of the synthetic oligonucleotides or building blocks. The melting temperature of 3WJ nanoparticle is significantly higher than that of the two-strand complexes, which is beneficial for future therapeutic applications at physiological temperatures.

We next tested the stability of the L-3WJ nanoparticles in human serum. Both L- and D-3WJ nanoparticles were incubated with 25% human serum for up to 24 hours and the presence or absence of degradation products were assessed by gel electrophoresis (Figure 1E). The band corresponding to the L-3WJ nanoparticle remained stable throughout all time points, while the native D-3WJ was completely degraded within 1 hour. These data indicate increased stability of the L-3WJ nanoparticles in human serum due to their altered chirality and lack of recognition by cellular RNAses.

### Effect of the L-3WJ nanoparticles on hematopoietic stem cells

Our objective is to use the L-RNA nanoparticle as potential therapeutics in disease treatment, therefore, it’s important to determine whether the nanoparticles cause unwanted toxicity to normal cells. We used human CD34^+^ cell as model to investigate whether the L-RNA nanoparticles affected growth and survival of human hematopoietic progenitor cells in culture. Progenitor cells were treated with different concentrations of L-RNAs (1-100 nM) or a vehicle (buffer) control for 14 days. L-RNA nanoparticles had no impact on formation of colonies (Figure 1F), indicating that the synthetic L-RNA nanoparticles, as an artificial biomaterial, don’t produce observable cellular toxicity to the human hematopoietic stem cells. The result suggests that L-RNA doesn’t present toxicity as the potential therapeutics.

### Binding and internalization of L-RNA nanoparticles mediated by folic acid

The ability to deliver drugs with precision and safety is a goal of nanoparticle technology. This can be achieved by regulating the delivery-dependent loading, targeting, and release of a specific drug, all while aiming to optimize its therapeutic potential. In order to test the L-3WJ nanoparticle as a precise drug delivery device, we synthesized a 5’-Cy5 fluorophore-labelled L-nanoparticle to monitor the binding and delivery process (L-3WJ_b-cy5_).

In addition, for the nanoparticle’s specific targeting cancer cells, we attempted to label the L-RNA with folic acid (FA), which is being actively explored in clinical trials and has shown potential in improving the precision and effectiveness of cancer therapies^31^. Folic acid, a vital B vitamin, has emerged as a promising agent for targeted drug delivery in cancer treatment. Its receptor, folate receptor alpha (FRα), is overexpressed in various cancer cells, including ovarian, breast, and lung cancers, while being minimally present in normal tissues^32^. This overexpression enables the selective targeting of cancer cells by conjugating folic acid to therapeutic agents, enhancing the delivery and efficacy of drugs directly to the tumor site while reducing systemic toxicity. Therefore, utilizing folic acid for L-RNA drug delivery is likely to leverage the natural cellular uptake mechanism, leading to improved cancer cell specificity and better treatment outcomes. To realize the FA-conjugation, the 5’-alkyne modified L-RNA was synthesized, using the 1-Ethynyl-dSpacer CE Phosphoramidite at the 5’-end. On the other hand, the folic acid-PEG-azido compound (Figure 2A) was chosen, which help to conjugate the FA group to RNA via click reaction^33^ (Figure 2B). Afterall, the multi-functional Cy5^+^, FA^+^ L-3WJ nanoparticle was prepared by annealing L-3WJ_a_, L-3WJ_b-cy5_ and L-3WJ_c-FA_ (Figure 2C), which was designed to detect the FA receptor biomarker expressed on cancer cell membrane.

**Figure 2.**
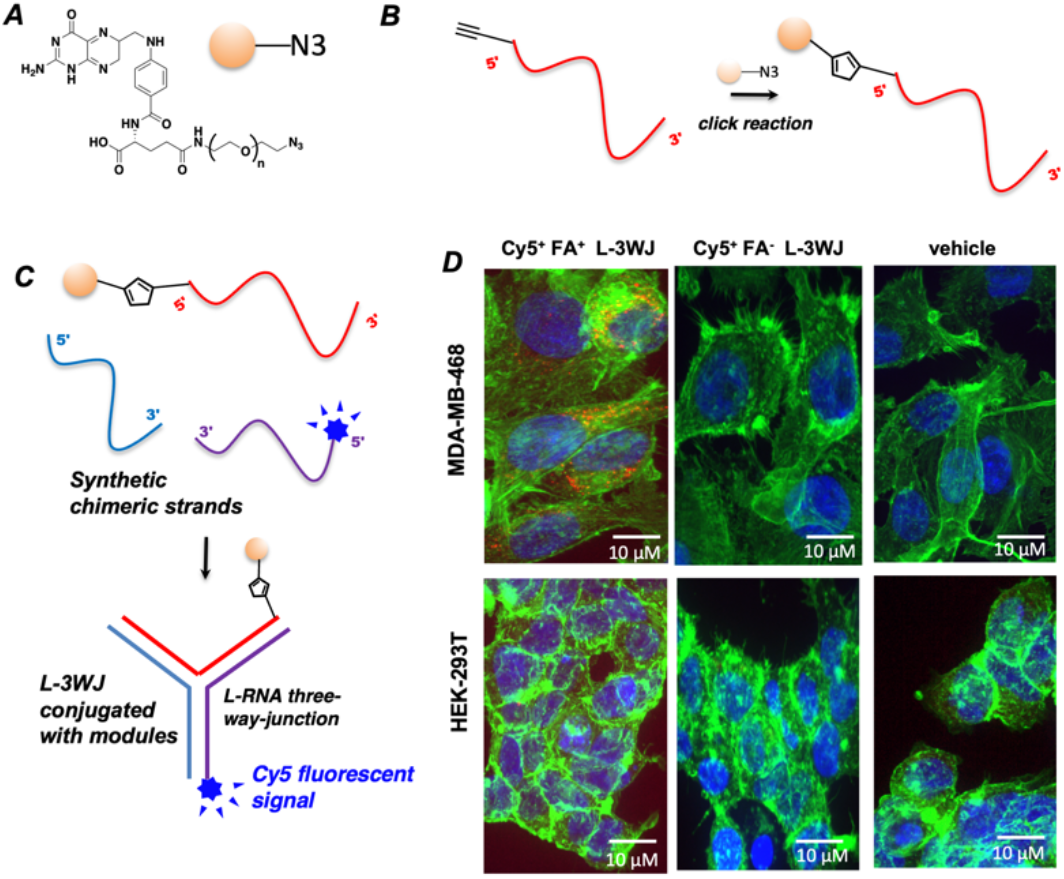
Binding and Internalization of L-RNA Nanoparticle Mediated by the FA group. (A) Chemical structure of FA-(PEG)_n_-azido compound used for labelling L-RNA. (B) Click reaction to label 5’-alkyne modified L-RNA. (C) Schematic presentation of construction of the Cy5^+^, FA^+^ L-3WJ, containing FA (orange ball) targeting FA receptor and Cy5 fluorophore (blue). (D) Confocal microscope images indicating the binding and internalization of L-3WJ into MDA-MB-468 cells (blue: nuclei; green: cytoskeleton; red: L-RNA nanoparticles, the concentration of each fluorescent RNA nanoparticles in treatment is 50 nM).

To facilitate the successful administration of anti-cancer agents into cancer cells, the RNA nanoparticle must first bind to the receptor on cancer cell membrane, and penetrate the cell membrane, allowing for the subsequent release of therapeutics into the cytoplasm. To validate the efficient delivery of L-RNA nanoparticle, the binding and internalization of RNA nanoparticles into cancer cells was examined by confocal microscopy.

Media containing 50 nM Cy5^+^, FA^+^ L-3WJ nanoparticle was used to treat the TNBC cell line MDA-MB-468 and the normal cell line HEK293t that does not naturally express folate receptors. For comparison, the same incubations were performed using the Cy5 labelled L-3WJ nanoparticle without the FA aptamer (Cy5+, FA-L-3WJ), to validate the specific targeting. The cell culture media for drug treatment and cell growth contained Dulbecco’s Modified Eagle Medium and Fetal Bovine Serum without heat inactivation, to test the efficiency of L-RNA. Following the nanoparticle incubation for 2h and washing steps, the cells were fixed, the nuclei were stained with DAPI, and the cytoskeletons were stained with FITC-conjugated phalloidin.

Confocal microscopy images show increased fluorescence from Cy5^+^, FA^+^ nanoparticles following incubation with MDA-MB-468 cells, but not with HEK-293T cells (Figure 2D). Furthermore, the observed Cy5 signals were surrounding the stained nuclei, suggesting the nanoparticles were internalized and delivered to cytoplasm of MDA-MB-468 cells. As the comparison, the Cy5^+^, FA^-^ L-3WJ were not binding or delivered into either cancer cells or normal cells, due to the lack of FA molecule to direct cell targeting. This successful detection of the L-RNA nanoparticle within cancer cells holds great promise for the potential to administer therapeutics with a high degree of stability and precision.

### Binding of doxorubicin to double stranded L-DNA

As the FDA-approved anti-cancer drug^34^, DOX binds to the complex formed by topoisomerase and double stranded DNA in human gene, stops the necessary unwinding of DNA, disrupts DNA synthesis and repair, and interferes with cancer cell growth^35^. However, its similar disruptive effects on healthy cells limit the maximum dose administered in the clinic^36^. In addition, Although DOX is water soluble, it forms a dimerized complex and precipitates when interacting with PBS buffer^37^. This insolubility in physiological conditions has hindered the clinical application of DOX in cancer treatment. As a result, there has been a number of studies focused on enhancing the pharmacokinetics of DOX through the implementation of different delivery systems ^38, 39^. Along these lines, using L-3WJ nanoparticles, we expect to take advantage of the excellent solubility of nucleic acid to stabilize DOX, thereby resulting in improved DOX delivery.

Native double-stranded DNA has been widely applied to conjugate with doxorubicin for drug delivery. The binding involves a well-understood mechanism where doxorubicin intercalates between the base pairs of DNA, facilitated by the planar aromatic rings of doxorubicin, which slide between the stacked bases of the DNA double helix^40^. This binding is also stabilized by van der Waals forces and hydrogen bonds between the doxorubicin molecule and the DNA bases. The primary amine on doxorubicin can also form ionic interactions with the phosphate backbone of DNA, further stabilizing the complex.

To use our mirror-image nucleic acid system to deliver chemotherapy, the small molecule drug needs to be stably loaded onto the mirror-image nucleic acid scaffold first. However, the binding of doxorubicin to L-DNA, the mirror image of natural D-DNA, raises questions due to differences in chirality. The difference in chirality means that the spatial arrangement of the bases and the overall helical structure are inverted in L-DNA compared to D-DNA. As a result, the intercalation and binding sites that doxorubicin typically recognizes in D-DNA might not be present or accessible in the same way in L-DNA. The chirality of the interacting molecules is crucial in molecular recognition processes, and many small molecules, including drugs, have a strong preference for binding to one enantiomer over the other. Given that doxorubicin has evolved to interact with D-DNA, its ability to bind to the mirror image L-DNA is uncertain and requires experimental verification. The unique spatial configuration of L-DNA might not provide the necessary alignment for effective intercalation and binding, thereby impacting the potential for L-DNA to serve as a carrier for doxorubicin delivery. This investigation will provide valuable insights into the potential use of L-DNA in drug delivery systems and could lead to the development of novel therapeutic strategies.

We first synthesized a double-stranded L-DNA, which contained a 16 nucleotides overhang, allowing Watson-Crick base pairing with another overhang of the L-3WJ nanoparticle. Given that DOX has been known to bind to D-DNA through hydrogen bonding with guanine, particularly in instances where another guanine is present on the opposing strand^41^, we designed several 5’-GC-3’/3’-CG-5’ sequences in dsDNA that can potentially serve as binding sites for DOX (Figure 3A). We measured the binding ratio of DOX/dsL-DNA by titrating DOX against various L-DNA concentrations and monitoring the corresponding fluorescent intensity from the DOX molecule. When using excitation at 480 nm and emission at 500-700 nm^42^, DOX solution presents a maximal emission near 600 nm (Figure 3B). When the concentration of dsL-DNA increases, the fluorescence intensity at 600 nm was gradually quenched due to the interaction between DOX and L-DNA. When the ratio of DOX/L-DNA reaches ∼5, the fluorescence was almost completely quenched. The findings reveal that approximately 5 molecules of doxorubicin bind to each dsL-DNA molecule, consistent with our DNA sequence design. To compare this result to DOX binding with native DNA, we then performed the same experiment using the synthetic D-DNA. The obtained result was similar to L-DNA/DOX binding. Under the same ratio, the DOX/L-DNA group presented slightly stronger fluorescent intensity than DOX/D-DNA group. Our results indicate that, doxorubicin binds to L-DNA, the mirror image of natural D-DNA, but the binding affinity is slightly weaker compared to its interaction with D-DNA. Our observation is similar to the reported exploration before, that mirror-image DNA tetrahedron can be used to load doxorubicin for specific tumor targeting^43^. The reduced binding affinity between L-DNA and DOX may be attributed to the differences in the chiral environment between L-DNA and D-DNA. In L-DNA, the altered spatial orientation and chiral configuration likely hinder the optimal intercalation, but still maintain the strong intercalation that are more favorable in double stranded DNA.

**Figure 3.**
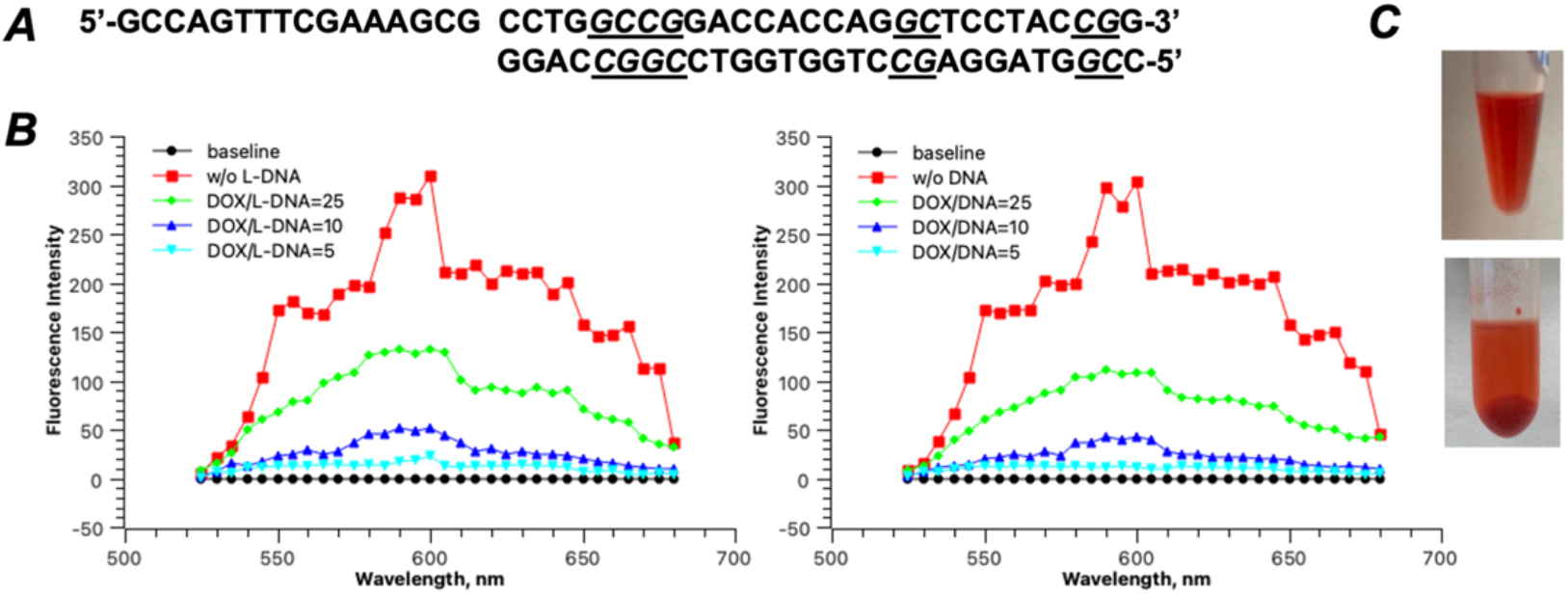
Conjugation of DOX to dsDNA and the solubility improvement. (A) The designed dsL-DNA sequence which can conjugate to L-3WJ and get intercalated by DOX. The underlined nucleotides indicate the GC rich region where DOX potentially binds. (B) Fluorescence characteristics of DOX in water before binding to dsL-DNA or dsDNA and after titrating by dsL-DNA or dsDNA. (C) Solubility comparison of DOX/dsL-DNA conjugate in PBS (top) buffe and DOX only in PBS buffer (bottom). The images show the visual precipitate of DOX drug in PBS buffer after heating at 37 °C.

By forming the conjugate of dsL-DNA/DOX, we can greatly enhance the solubility of DOX in physiological buffer. DOX hydrochloride was dissolved in 1x PBS buffer (20 mM phosphate, 300 mM NaCl, pH 7.2) to yield concentration of 5 mg/mL. By incubating the solution at 37 °C for 10 min, a red colored precipitate was observed, probably due to the formation of covalently bonded DOX dimer (Figure 3C, top). In comparison, the solution of dsL-DNA/DOX conjugate in a PBS buffer was completely dissolved at the same DOX concentration (Figure 3C, bottom). Even after incubating at 37 °C for 12 h, the complex solution showed no visible precipitate. In addition to the improvement in solubility of DOX in PBS buffer, the drug loading capacity can be enhanced by using our L-DNA design. In the practical terms, the L-DNA/DOX conjugate can further assemble with L-3WJ nanoparticle, and more DOX molecules can be incorporated into one nanoparticle by optimizing the DNA sequence.

### Use L-3WJ nanoparticle to delivery siRNA for gene silencing

Combining siRNA delivery with chemotherapy offers a promising strategy to combat cancer treatment challenges, particularly drug resistance^44^. Chemotherapy often leads to the development of resistance in cancer cells, making subsequent treatments less effective and contributing to cancer recurrence. By co-delivering siRNA, we aim to target and regulate specific genes associated with drug resistance mechanisms^45^. This dual approach can help to sensitize cancer cells to chemotherapy, thereby enhancing its efficacy and preventing the re-emergence of resistant cancer cells. This synergistic method holds potential for improving treatment outcomes and achieving more durable remissions in cancer patients.

To achieve this dual therapeutic approach, we propose using our L-RNA nanoparticles to deliver siRNA. L-RNA, the mirror image of natural D-RNA, offers enhanced stability as a significant advantage in the bloodstream and cellular environments. With the conjugate of folic acid, these nanoparticles can efficiently target cancer cells and deliver the therapeutic agents via endocytosis. However, there are concerns to address, particularly whether the RNA interference (RNAi) machinery, such as Dicer, will recognize and process the L-RNA-siRNA conjugate effectively. Dicer’s ability to cleave and activate siRNA is crucial for the subsequent gene silencing mechanism^46^. If Dicer does not recognize or properly process the L-RNA-siRNA due to the chirality compatibility issue, the intended gene regulation will not occur, rendering the therapeutic approach ineffective. Another critical concern is whether the entire L-RNA-siRNA conjugate can escape from the endosome after cellular uptake. Efficient endosomal escape is necessary for the siRNA to reach the cytoplasm and engage the RNA-induced silencing complex (RISC) to function effectively^47^. If the L-RNA nanoparticles fail to escape the endosome, the siRNA will be trapped and degraded, preventing it from silencing the target genes associated with drug resistance. Ensuring that our L-RNA-siRNA conjugates are designed to facilitate endosomal escape, alongside their compatibility with the natural RNAi pathway, is essential for achieving the desired therapeutic outcomes. Additionally, potential issues such as immune recognition, cellular uptake efficiency, and the safe release of siRNA within target cells must also be carefully addressed.

To evaluate the efficacy of our L-RNA nanoparticle delivery system, we tested it using siRNA targeting the MCL1 gene. MCL1 (Myeloid Cell Leukemia 1) is an anti-apoptotic member of the Bcl-2 family of proteins, playing a crucial role in cell survival and resistance to apoptosis^48^. Overexpression of MCL1 is commonly observed in various cancers and is associated with increased tumor cell survival, resistance to chemotherapy, and poor clinical outcomes. By targeting MCL1 with siRNA, we aim to downregulate its expression, thereby promoting apoptosis in cancer cells and sensitizing them to chemotherapeutic agents. This makes MCL1 a critical target for cancer therapy, as its inhibition can significantly enhance the effectiveness of treatments and potentially overcome drug resistance in cancer cells.

Therefore, in our study, we aimed to investigate the delivery effectiveness of siRNA in silencing the MCL1 gene. We designed specific siRNA sequences (Figure 4A) to target the MCL1 gene and employed Western blot analysis to quantitatively assess the gene silencing effects. To evaluate the impact of folic acid modification on L-RNA nanoparticle delivery, we prepared L-RNA nanoparticles conjugated with varying amounts of folic acid groups to evaluate the impact of folic acid density on delivery efficiency. In parallel, we used lipofectamine, a widely used transfection reagent, as a benchmark to compare the gene silencing efficacy of our L-RNA nanoparticle delivery system. MDA-MB-468 cells, which overexpress the folate receptor, were cultured in DMEM medium with 10% FBS. For drug delivery assay, cells were seeded in 6-well plates and incubated to reach ∼80% confluency. The nanoparticle conjugates were diluted in Opti-MEM medium and added to the cells at a final concentration of 1 µM. Cells were incubated with the transfection mixture for 4 hours, after which the medium was replaced with fresh medium. For comparison, lipofectamine 3000 was used according to the manufacturer’s protocol. Cells were treated with different lipofectamine-L-RNA conjugate complexes also at a final concentration of 1 µM. After 48 hours of transfection, cells were harvested and lysed in RIPA buffer to release the proteins. Equal amounts of protein were separated by SDS-PAGE and transferred to PVDF membranes. Membranes were blocked with 5% non-fat dry milk in TBST and probed with primary antibodies against MCL1 and β-actin (as a loading control) overnight at 4°C. The peroxidase-conjugated secondary antibody was used for chemiluminescence analysis.

**Figure 4.**
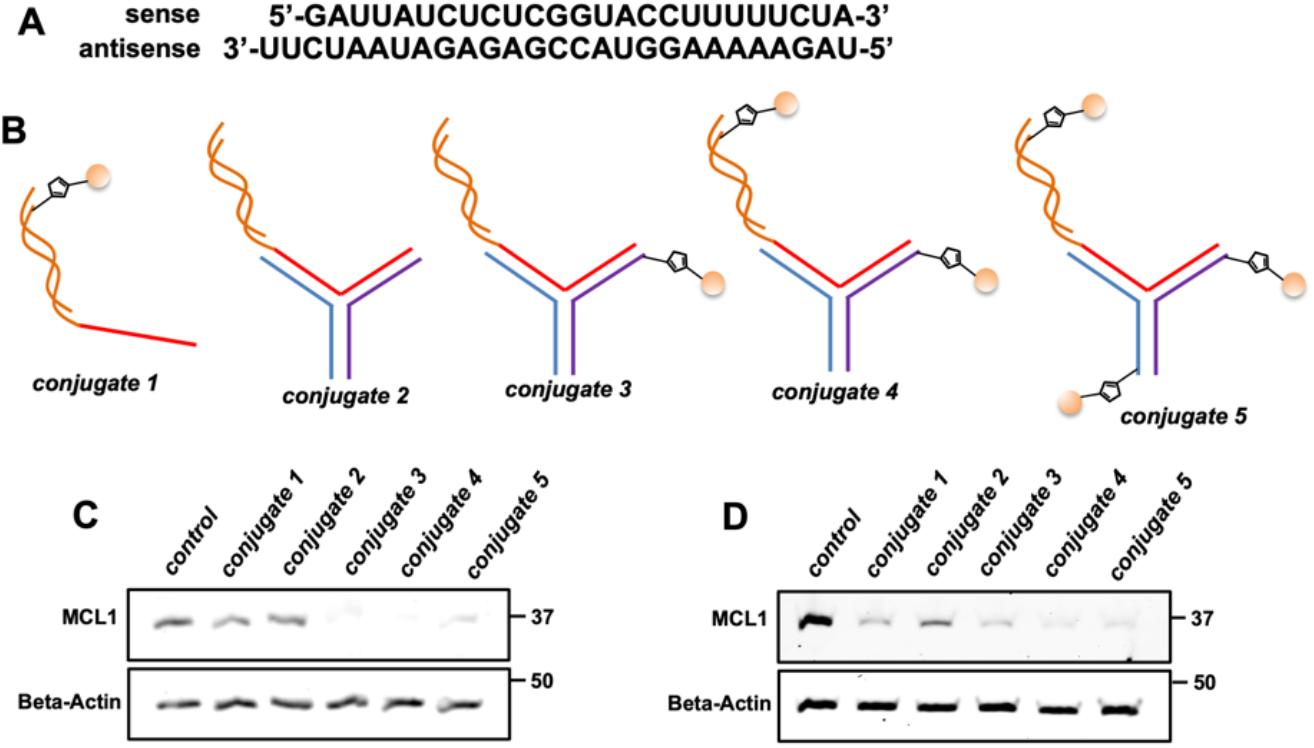
Gene silencing effects by using different nanoparticle conjugates to treat MDA-MB-468 and HEK293t cells. (A) Sense and antisense sequences of siRNA to target MCL1 gene. (B) Schematic representations of different nanoparticle designs. Orange ball: FA group; yellow curve: MCL1 siRNA. (C-D) Western blot analysis of suppressing MCL1 protein expression with different nanoparticles, by FA-receptor mediated endocytosis on MDA-MB-468 cells (C) and Lipofectamine 3000 transfection on MDA-MB-468 cells (D). Gels images depicted are representative of triplicate experiments.

Western blot results were shown in Figure 4C and 4D. In the FA-dependent delivery, no transfection agent was used (Figure 4C). When the FA-conjugated double stranded siRNA was used (conjugate 1), MCL1 gene silencing was observed, and the protein expression was suppressed by ∼40%. In the group of L-RNA-siRNA conjugate 2 without FA labeling, no gene silencing was observed. In comparison, when the L-RNA-siRNA conjugates were labeled by FA at different positions (conjugate 3, 4 and 5), significant gene silencing was generated. In general, the L-3WJ complexes containing two FA moieties (conjugate 4) and one FA moiety (conjugate 3) lead to slightly stronger gene silencing than the conjugate 5, which contained three FA groups (∼90% suppression over ∼80% suppression). This observation agrees with other lab’s report that, folic acid facilitates the selective delivery of DNA tetrahedron nanoparticle into cells, and the FA groups on DNA nanoparticle are critical for the optimal endosomal escape^49^. Surprisingly, conjugate 5 containing three FA groups didn’t cause the strongest gene silencing effect. Theoretically, more FA conjugation can help the nanoparticle to have better binding affinity to cancer cells, and contribute more to the endosomal escape of the nucleic acid agents. However, we speculate that conjugating folic acid to our L-3WJ might also introduce steric hindrance, which can affect the binding affinity of the nanoparticles to their target receptors. This could reduce the efficiency of overall cellular uptake and gene silencing. Plus, the chemical modification process used to attach folic acid to the RNA nanoparticles could potentially alter the stability of the nanoparticles. This could lead to unwanted aggregation of the nanoparticles. All these factors generated the experimental result we observed. The relationship between numbers of folic acid groups and size of L-RNA nanoparticle will needs in-depth investigation in the future.

As comparison, when delivering the nucleic acid drugs using lipofectamine 3000, all the conjugates presented prominent gene silencing effects (Figure 4D). Interestingly, the conjugates containing folic acid groups (conjugate 1, 3, 4, 5) showed slightly stronger gene silencing effects (over 95%) compared to conjugate 2 without folic acid (∼85%). One possible explanation for this observation is the folic acid molecules facilitate the formation of more stable lipid-RNA complexes with Lipofectamine 3000, improving the overall efficiency of the transfection process. Additionally, the stronger silencing effect is also possibly the combination of Lipofectamine 3000-dependent delivery and FA-receptor dependent endocytosis, in which more FA molecules lead to more effective uptake of the nanoparticles by the cells.

In conclusion, our experimental results indicated that the folic acid-conjugated L-RNA nanoparticles successfully delivered siRNA and achieved gene silencing of MCL1, as evidenced by the reduced protein levels in the Western blot analysis. The conjugated FA molecules not only help the nanoparticles to target the cancer cells but also contribute to the endosomal escape of the RNA drugs. Considering the overexpression of FA receptor on cancer cell membrane instead of normal cells, our L-RNA nanoparticle system is possibly a more precise and efficient strategy to delivery siRNA therapy for cancer treatment without interfering normal cells.

### Delivery of L-RNA nanoparticles to cancer cells for viability assay

After conjugating to L-DNA, DOX can be stabilized in solution to avoid precipitation. We then performed the dose-dependent Alamar Blue assay in both cancer and normal cell lines by using different nanoparticle conjugates (Figure 5A), to assess their therapeutic effects. The Alamar Blue assay is a widely used and reliable method for assessing cell viability and proliferation^50^. This colorimetric assay is based on the reduction of resazurin, a non-toxic and cell-permeable dye, to resorufin, a fluorescent and brightly colored compound, by metabolically active cells. The extent of this reduction, which can be quantified by measuring the fluorescence or absorbance, directly correlates with the number of viable cells. In the context of evaluating RNA nanoparticles’ drug delivery efficiency, the Alamar Blue assay provides us an effective means to determine the cytotoxicity and biocompatibility of these nanoparticles. The corresponding results were presented in Figure 5B. The x-axis represents the concentration of the therapies, with each sample subjected to serial dilutions by a factor of two, starting from 500 nM. The y-axis indicates the relative cell viability compared to the control group, where no drug was added.

**Figure 5.**
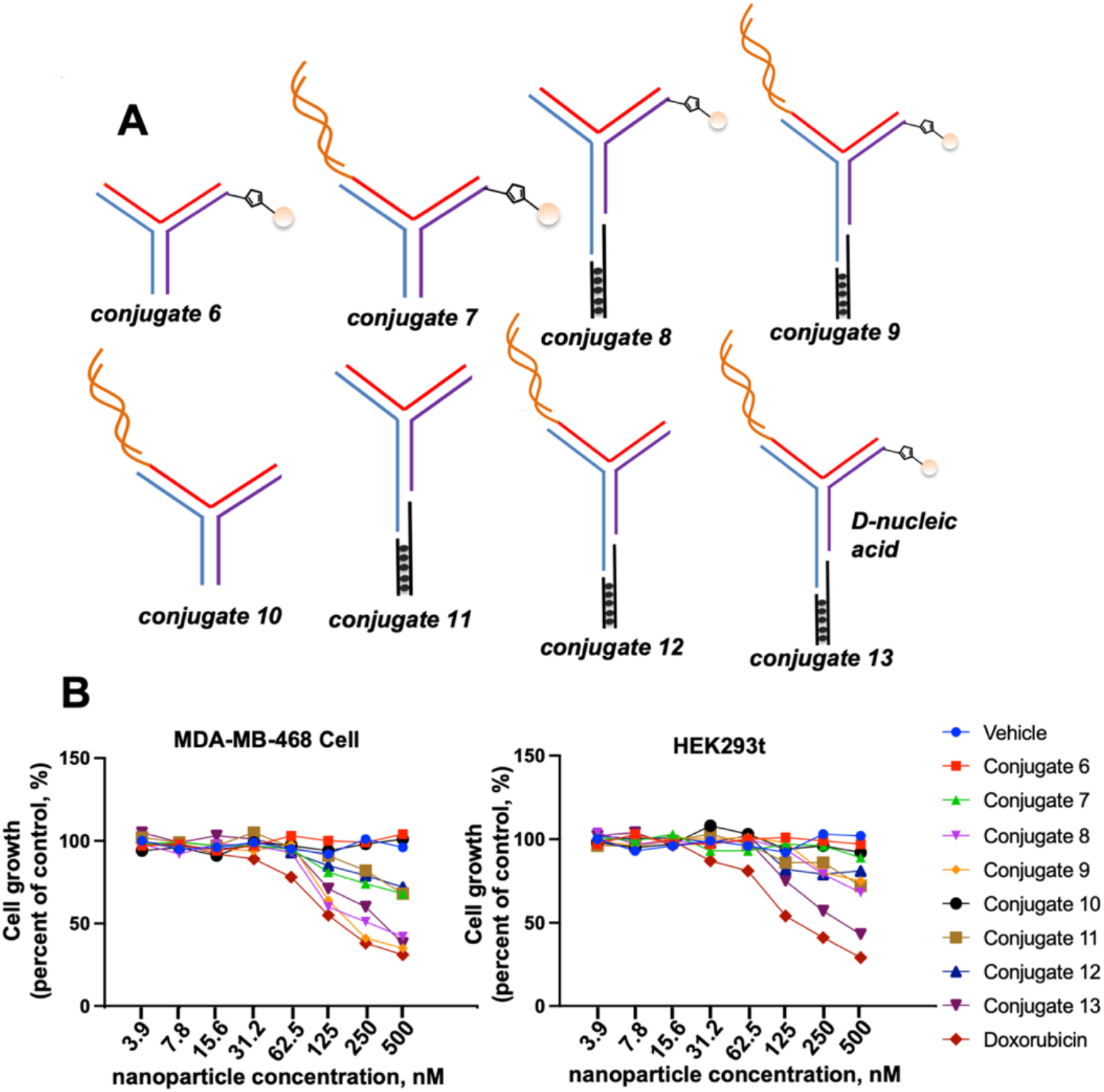
Inhibition of cell growth in MDA-MB-468 and HEK293t cells by the therapeutic nanoparticles. (A) Schematic representations of different nanoparticle designs. Black dots: DOX; orange ball: FA group; yellow curve: MCL1 siRNA. (B) Alamar blue assay using different nanoparticles to treat MDA-MB-468 and HEK293t cells. The cell growth inhibitory effects were compared between different nanoparticle conjugates. Data are representative of triplicate experiments.

We first performed the Alamar Blue assay in MDA-MB-468 cells. Among all the therapeutic agents, the most potent were the conjugates containing both a folic acid group and doxorubicin (conjugates 8, 9, and 13). These three nanoparticles suppressed cancer cell growth by over 50% at the 500 nM concentration treatment, demonstrating the same inhibitory capacity as free doxorubicin. Interestingly, conjugate 9 showed slightly stronger toxicity than conjugate 8, likely due to the MCL1 siRNA’s role in regulating cancer cell apoptosis. Nanoparticles carrying doxorubicin without folic acid conjugation (conjugates 11 and 12) also exhibited toxicity. At a 500 nM concentration, cancer cell growth was inhibited by approximately 30%, which we speculate is due to doxorubicin molecules dissociating from the L-DNA and passively penetrating the cell membrane to induce cell death. Treatment with conjugate 7, which contains FA and MCL1 siRNA, also resulted in ∼30% cell growth inhibition, likely from the promotion of apoptosis by targeting MCL1 with siRNA, thereby reducing MCL protein expression. When the therapeutics contained only the folic acid group (conjugate 6) or siRNA (conjugate 10), no toxicity was observed, and cancer cell growth remained as robust as in the vehicle group.

We then performed the same experiment in HEK293t cells, which do not express the folic acid receptor on their cell membranes. Conjugates 6, 7, and 10 did not exhibit observable toxicity, and cell viability was identical to the vehicle group. This is because either no doxorubicin was conjugated to the nanoparticle, or the FA could not direct the drug delivery due to the lack of FA-receptor interaction. Similar to the results in MDA-MB-468 cells, conjugates 11 and 12 showed modest toxicity, inhibiting cell proliferation by approximately 32%. This is likely due to doxorubicin leaking into the media during incubation. Additionally, the toxicity of conjugates 8 and 9 was significantly reduced when treating HEK293t cells, with about 35% cell growth inhibition observed, similar to conjugates 11 and 12, which lack the FA group. Compared to the toxicity in cancer cells (over 50% inhibition), the FA group in conjugates 8 and 9 did not facilitate efficient endocytosis to deliver DOX, indicating that the observed toxicity was still due to the passive penetration of DOX in the cell culture media. Furthermore, conjugate 13, the D-RNA nanoparticle conjugate, did not show optimal selectivity of cells in the viability assay. In both cancer and normal cell groups, conjugate 13 exhibited strong cell growth inhibition (over 50%). Given the lack of FA-receptor mediated endocytosis to deliver DOX in HEK293t cells, we speculate that the significant toxicity is due to the high concentration of DOX in the media. When D-nanoparticles were included during drug treatment, the serum in the cell culture media caused D-DNA degradation, leading to the complete release of DOX molecules from the conjugates and significant passive penetration of the cell membrane, resulting in cell death. The different inhibitory effects of conjugate 9, 12 and 13 at 500 nM treatment are shown in Figure 6.

**Figure 6.**
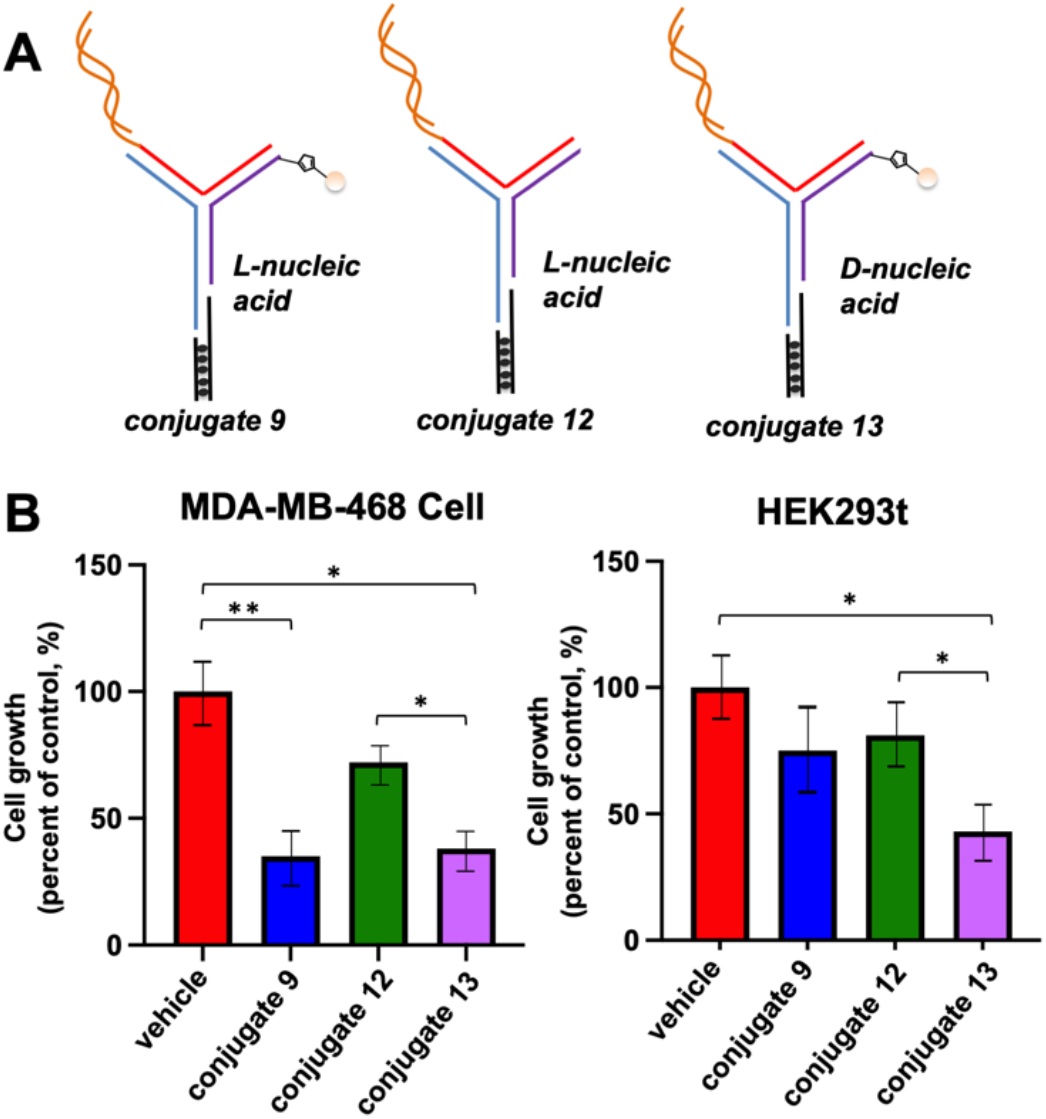
Comparison of inhibition of cell growth in MDA-MB-468 and HEK293t cells by three nanoparticles. (A) Schematic representations of conjugate 9, 12 and 13. Black dots: DOX; orange ball: FA group; yellow curve: MCL1 siRNA. (B) Alamar blue assay using the three different nanoparticles to treat MDA-MB-468 and HEK293t cells. L-nucleic acid-based conjugate 9 shows different inhibitory effects between MDA-MB-468 and HEK293t cells, while the other two conjugates can’t differentiate MDA-MB-468 and HEK293t cells. Error bars represent the mean ± SEM from three independent experiments (n = 3). Statistical significance was determined using an unpaired two-sided t-test. Significance symbols: *P < 0.05, **P < 0.01.

In summary, our L-RNA-based nanoparticle conjugate, incorporating both folic acid (FA) groups and doxorubicin (DOX), demonstrates selective targeting for cancer cells and exhibits effective inhibition of cancer cell growth. The FA-directed drug delivery system enhances the inhibitory effects on cancer cells while minimizing interference with normal cell proliferation. Traditionally, DOX is released from DNA carriers via nuclease digestion of double-stranded DNA (dsDNA) within cells during endocytosis. However, L-DNA is not recognized by nucleases, raising questions about the release mechanism of doxorubicin within cancer cells. Possible mechanisms include: 1) the acidic environment within endosomes or lysosomes could induce structural changes in L-DNA, leading to the release of doxorubicin; and 2) cellular proteins and other molecules might displace doxorubicin from L-DNA, allowing it to intercalate into the genomic DNA of cancer cells. Detailed investigations into these mechanisms are necessary for a comprehensive understanding. Additionally, the FA group enhances the delivery of siRNA, thereby improving the combined therapeutic effect. Compared to D-RNA nanoparticles, the L-RNA system offers a more effective and safer strategy for delivering multiple therapeutics. This approach holds promise for improved targeting and efficacy in cancer treatment.

## CONCLUSION

In this study, we have successfully demonstrated the construction and application of a novel L-RNA-based nanostructure to enhance drug delivery efficiency and stability. Our findings indicate that the L-RNA three-way junction (L-3WJ) nanoparticles exhibit remarkable stability in human serum, significantly outperforming traditional D-RNA-based systems. The targeted delivery of these nanoparticles was facilitated by folic acid functionalization, which enhanced their uptake by cancer cells overexpressing the folate receptor, leading to improved cytotoxicity. Notably, we discovered that folic acid not only aids in targeted delivery but also significantly improves the intracellular escape of siRNA, thereby enhancing the effectiveness of siRNA-mediated gene silencing. The conjugation of doxorubicin and siRNA to the L-RNA nanoparticles resulted in significant inhibition of cancer cell growth, with the combination of doxorubicin and MCL1 siRNA showing synergistic effects that further enhanced therapeutic outcomes.

While our results underscore the potential of L-RNA nanoparticles as a robust platform for targeted cancer therapy, there are several avenues for future research. Optimizing the delivery efficiency and therapeutic efficacy of these nanoparticles is crucial, and exploring additional targeting ligands could improve specificity. In vivo studies are essential to validate our findings and assess the biocompatibility and overall efficacy of L-RNA nanoparticles in animal models. Despite their enhanced stability, potential challenges include efficient and scalable synthesis, immune responses, and achieving consistent and controlled drug release, which will be our future direction. To address these limitations, strategies such as incorporating cell-penetrating peptides to improve intracellular delivery, surface modifications like PEGylation to reduce immunogenicity, and developing stimuli-responsive systems for controlled release may need to be considered. Additionally, advancing synthesis techniques and purification processes will be pivotal for the practical application of L-RNA nanoparticles in clinical settings.

In conclusion, L-RNA-based nanostructures offer a promising and versatile platform for targeted and efficient drug delivery, with the potential to revolutionize cancer therapy and other medical treatments. The enhanced stability, targeted delivery, and improved intracellular escape of siRNA facilitated by folic acid functionalization represent significant advancements in the field. Continued research and development will be essential to fully realize their capabilities and address existing challenges, paving the way for next-generation therapeutic strategies.

## SUPPLEMENTARY DATA

Supplementary Data are available at xxxx.

## ACKNOWLEDGEMENTS

We thank Dr. Zhang and members for helpful discussions, insightful commentary, and careful revision of the manuscript. We thank Dr. Kirk Staschke and Dr. Ricardo Cordova for their valuable assistance and helpful discussions during the experiments. We thank Dr. J. Meng and the Chemical Genomics Core at IUSM for spectropolarimeter analysis.

