## supporting information for "Construction of mirror-image RNA nanostructure for enhanced biostability and drug delivery efficiency"

#### CONTENTS

1. General methods S2

2. List of sequences of all the oligonucleotides S3

3. Protocol of labeling RNA with folic acid molecule by Click reaction S3

4. Inhibitory effect comparison. S4

**1. General Methods.**

**1a. General considerations.** All chemicals were purchased from Fisher Scientific, VWR or Sigma-Aldrich unless otherwise noted. All the nucleoside phosphoramidites for solid-phase synthesis were from Chemgenes Corporation or Glen Research. All oligonucleotides, including fluorophore modified RNAs, 2'-F-modified RNAs, alkyne-modified RNAs and RNA-DNA chimeric strands, were synthesized in house. The alkyne modified phosphoramidintes were purchased from Hongene Biotech.

**1b. RNA oligonucleotides synthesis.** RNA oligonucleotides were synthesized by standard solid-phase phosphoramidite chemistry on a ABI RNA/DNA oligonucleotide synthesizer. Cleavage and elution of the full-length products from 1 μmol universal CPG-solid support columns, as well as removal of protecting groups on the nucleobases and phosphates, was carried out by heating the solid support for 15 min at 65 °C. The resultant homogeneous mixtures were first concentrated under reduced pressure for 2 hrs on a Genevac EZ-2 table top speedvac system (Genevac, Stone Ridge, NY), then lyophilized to dryness on a Labconco Benchtop 4.5 L freeze-drier (Labconco, Kansas City, MO) at <200 mTorr overnight to afford off-white solid residues. The residues were then resuspended in 115 μL of DMSO. 75 μL of TEA and 65 μL of TEA·3HF were added, and the solutions were heated for 2.5 hr at 65 °C to remove the TBDMS protecting group on the ribose 2'-hydroxyl group. The mixtures were homogeneous and pale to golden yellow in color. After cooling to room temperature (~30 mins), the RNA sample was desalted and detritylated by using the Glen-Pak RNA purification cartridge, following the protocol from Glen Research Inc. The purification of the desired products was performed by running denaturing polyacrylamide gels (urea-PAGE). The gel pieces containing the RNA fractions were collected, fragmented, and soaked in water for 12 hours. The RNA samples were then desalted by RNA precipitation using sodium chloride and ethanol. The precipitates were spun down (12000 rpm, 40 mins) and supernatants were removed by decanting. The resulting white solids were washed twice with absolute ethanol; the samples were then dried under high vacuum overnight before dissolving into water for appropriate concentrations.

#### 2. List of sequences of all the oligonucleotides

| entry | Sequence (5′ →3′) | nanoparticle |
|---|---|---|
| L-3WJ$_a$ | L-rUUGCCAUGUGUAUGUGGG | L-3WJ |
| L-3WJ$_b$ | L-rGGAUCAAUCAUGGCAA | |
| L-3WJ$_c$ | L-rCCCACAUACUUUGUUGAUCC | |
| D-3WJ$_a$ | D-rUUGCCAUGUGUAUGUGGG | D-3WJ |
| D-3WJ$_b$ | D-rGGAUCAAUCAUGGCAA | |
| D-3WJ$_c$ | D-rCCCACAUACUUUGUUGAUCC | |
| Cy5-L-3WJ$_b$ | Cy5-L-rGGAUCAAUCAUGGCAA | FA$^+$/Cy5$^+$ L-3WJ |
| FA-L-3WJ$_c$ | FA-L-rCCCACAUACUUUGUUGAUCC | |
| DOX-L-3WJ$_b$ | L-dCGCTTTCGAAACTGGC<br>-rGGAUCAAUCAUGGCAA | FA$^+$/DOX$^+$ L-3WJ |
| dsL-DNA-1 | L-dGCCAGTTTCGAAAGCGCCTGGCC<br>GGACCACCAGGCTCCTACCGG | |
| dsL-DNA-2 | L-dCCGGTAGGAGCCTGGTGGTCCGGCCAGG | |
| L-3WJ$_a$-siRNA | rGAUUAUCUCUCGGUACCUUUUUCUA-spacer-<br>L-rUUGCCAUGUGUAUGUGGG | siRNA$^+$/FA$^+$ L-3WJ |
| siRNA-<br>antisense | rUAGAAAAAGGUACCGAGAGAUAAUCUU | |
| FA-L-3WJ$_b$ | FA-L-rGGAUCAAUCAUGGCAA | |
| FA-L-3WJ$_a$-<br>siRNA | FA-rGAUUAUCUCUCGGUACCUUUUUCUA-<br>spacer-L-rUUGCCAUGUGUAUGUGGG | |

### underlined nucleotides represent the 2′-fluoro-pyrimidines.

#### 3. Label RNA with folic acid molecule by Click reaction

Incubate 40 mM $CuSO_4$ with 80 mM tris(3-hydroxypropyltriazolylmethyl)amine (THPTA) ligand in a 1:2 ratio for 5 minutes before the reaction. This solution is stable for several weeks when frozen. To the 50 uM alkyne-RNA solution 100 uL, add 10 uL 500mM Na-phosphate buffer (pH 7), then add an excess of 5 mM folic acid-PEG-azide solution 50 uL (4-50 eq). 25 equivalents of THPTA/$CuSO_4$ solution was added, followed by adding 2 uL 100 mM sodium ascorbate solution. The reaction stands at room temperature for 30 minutes. Folic acid labeled RNA strands were ethanol precipitated and quantified before nanoparticle assembly.

4. Inhibitory effect comparison of conjugates 9, 11 13 and doxorubicin in MDA-MB-468 and HEK293t cells.

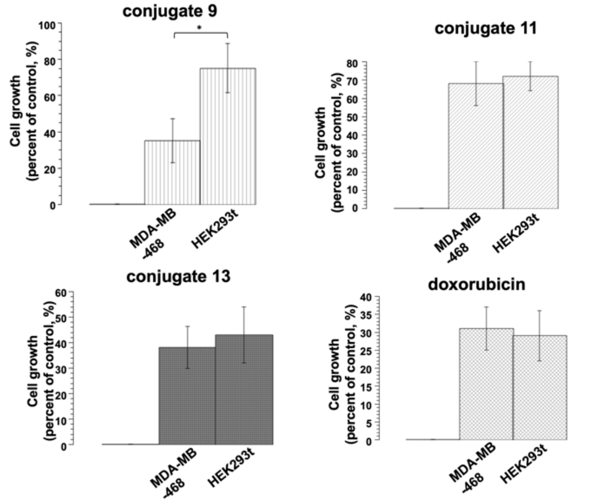


Figure S1. Alamar blue assay to show 4 different nanoparticles' therapeutic effects at 500 nM concentration treatment. The cell growth inhibitory effects were compared between cancer cell and normal cell. Only conjugate 9, which is an L-RNA nanoparticle conjugated with both FA and DOX, shows different inhibitory effects in MDA-MB-468 and HEK293t cells. Other nanoparticle conjugates can't specifically recognize different cell lines. Error bars represent the mean ± SEM from three independent experiments (n = 3). Statistical significance was determined using an unpaired two-sided t-test. Significance symbols: *P < 0.05.
